# Genetic diversity of Toscana virus glycoproteins affects the kinetics of virus entry and the infectivity of newly produced virions

**DOI:** 10.1101/2024.07.22.604564

**Authors:** Adrien Thiesson, Marie-Pierre Confort, Sophie Desloire, Alain Kohl, Frédérick Arnaud, Maxime Ratinier

## Abstract

Toscana virus (TOSV) is a pathogenic and transmissible *Phlebovirus* of the *Bunyavirales* order. Although TOSV is considered one of the leading causes of meningitis and encephalitis in humans during summer in the Mediterranean basin, its biology remains poorly characterized and neglected due to lack of tools to study the virus. To date, two principal genetic lineages (A and B) have been identified among TOSV-isolated strains based on phylogenetic analysis. The impact of TOSV genetic diversity on its biology is still unknown but highly relevant because it may influence the severity of the disease, viral tropism, and vaccine design. To address these questions, a reverse genetic approach based on two TOSV strains belonging to lineage A or B (i.e., TOSV-A and TOSV-B) and displaying different *in vitro* replicative fitness was used. Our results demonstrate that TOSV-A and TOSV-B have different Gn and Gc glycoproteins sequences which are responsible for the observed differences in terms of replicative fitness. Moreover, our data show that TOSV-A and TOSV-B display different entry kinetics and that newly-produced virions have different infectivity. This comparative approach allowed us to demonstrate that the genetic diversity of TOSV can significantly impact viral properties. This study highlights the need for a better molecular characterisation of the genome of circulating TOSV strains and, more specifically, of the viral Gn and Gc glycoproteins. Indeed, these proteins may strongly modulate viral pathogenicity and disease. Further work in this direction will provide important data to develop preventive strategies against this emerging pathogen taking into account TOSV glycoproteins genetic diversity.

**Authors Summary:** Toscana virus (TOSV) is a leading cause of aseptic brain infection in the Mediterranean basin during the summer. Despite the significant burden that TOSV represents to human health, the biology of this pathogen remains poorly understood and neglected. While distinct TOSV genetic lineages have been identified, the relationship between their genetic diversity and pathogenicity is still unclear. This point, however, is critical to understand the disease and design preventive strategies such as vaccines targeting circulating TOSV strains. Here, a reverse genetic approach was used to produce reassortant and chimeric viruses between two TOSV strains (referred to as TOSV-A and TOSV-B) belonging to the two main genetic lineages and displaying differential *in vitro* replication capacities. Our results show that the viral glycoproteins are key determinants in modulating TOSV replication. In addition, they demonstrate that viral entry and infectious viral particles production differ between TOSV-A and TOSV-B. This study provides the first evidence of differences in replication capacity between two genetically distinct TOSV viruses, and highlights the need for better molecular characterisation of circulating TOSV strains.

## Introduction

Toscana virus (TOSV) is an arbovirus of the *Bunyavirales* order and the *Phenuiviridae* family. It is transmitted by sand flies, mostly *Phlebotomus perniciosus* and *P. perfiliewi* [1]. Isolated for the first time in Italy in 1971 [2], TOSV is now found in 23 different countries around the Mediterranean basin [3–5]. Infection in humans is generally asymptomatic or leads to mild symptoms [6] but can progress to severe neurological disease following viral invasion of the central nervous system (CNS) [7,8]. TOSV is now considered as one of the leading causes of meningitis and encephalitis in humans during summer [9].

Two main distinct TOSV genetic lineages have been identified thus far based on phylogenetic analyses, and referred to as lineages A and B [3]. More recently, a third lineage (referred to as lineage C) was described based on partial viral sequences; however, this virus has not been isolated so far [10,11]. Despite the significant impact of TOSV on human health as an emerging cause of CNS infections, this virus remains an understudied pathogen. For instance, the impact of TOSV genetic diversity on viral pathogenicity is still unknown. Yet this represents an important point to address to understand TOSV disease and design preventive measures. Recently, we and others have developed reverse genetic systems for TOSV that allow manipulation of the viral genome [12,13], providing new opportunities to investigate TOSV molecular biology. In this study, two TOSV strains from lineages A (TOSV-A) and B (TOSV-B) were studied using reverse genetic approaches and reassortant viruses. Our results demonstrate that TOSV-A replicates at a higher rate than TOSV-B, due to differences in Gn and Gc coding sequences. In addition, our data show that TOSV-A and TOSV-B display different entry kinetics and that newly produced virions have different infectivity. Overall, our results highlight the significant impact of the genetic diversity of the glycoproteins of TOSV strains on virus biological characteristics. This points to the urgent need for more and wider molecular characterisation of circulating TOSV strains for a better understanding of this pathogen.

## Results

### Development of a reverse genetic system for TOSV-B

We previously published a reverse genetic system for one strain of TOSV lineage A TOSV (TOSV 1500590, referred to as wild-type [wt] TOSV-A) [12,14]. Here, we developed a reverse genetic system for one strain of TOSV lineage B (TOSV MRS2010-4319501, referred to as wt TOSV-B) [15]. To this end, the three antigenomic segments (Seg-S, Seg-M, and Seg-L) of the wt TOSV-B were amplified by RT-PCR and cloned into reverse genetic plasmids. Next, the sequences from these plasmids were compared to those of the wt TOSV-B strain reference genome available on GenBank [15]. Several substitutions were identified in all the segments: three in Seg-S, two in Seg-M, and seven in Seg-L (Fig 1A). The three substitutions in the Seg-S are in the coding sequence of the N protein: C521A and C522G, both resulting into a change of amino acid (His162Asp), and C524U (silent substitution). The two substitutions in the Seg-M are in the anti-genomic untranslated regions (UTRs): U80C, in the 5’ UTR, and G4096A, in the 3’ UTR. Finally, the seven substitutions in Seg-L are in the L polymerase coding sequence: G1071U (Asp785Glu), C2373A, G2379C, A2382C, A2388U, C4118U (Thr1367Ile), and G5224A (Val1736Met). To ensure that these substitutions were not generated during TOSV genome amplification, segments were again amplified by RT-PCR from wt TOSV B and re-sequenced. This confirmed that the cloned sequences were correct. Furthermore, these sequences were compared to those of other TOSV strains belonging to lineage B and available on GenBank. As shown in Fig 1B, these sequences are identical, except for substitution U80C in the anti-genomic 5’ UTR. Thus, the 5’ UTR and 3’ UTRs of each segment from our viral stock were re-sequenced by RACE PCR, confirming the U80C substitution in the wt TOSV-B antigenome. In addition, when comparing the results from the RACE-PCR and the sequences of the TOSV-B plasmids, two additional nucleotide substitutions were identified: one in the Seg-S 3’ UTR (C1860A), and the other one in the Seg-L 5’ UTR (G6A). These 2 substitutions were modified by site-directed mutagenesis to correct the sequence of the two plasmids containing the Seg-S and Seg-L for reverse genetics purposes.

**Fig 1.**
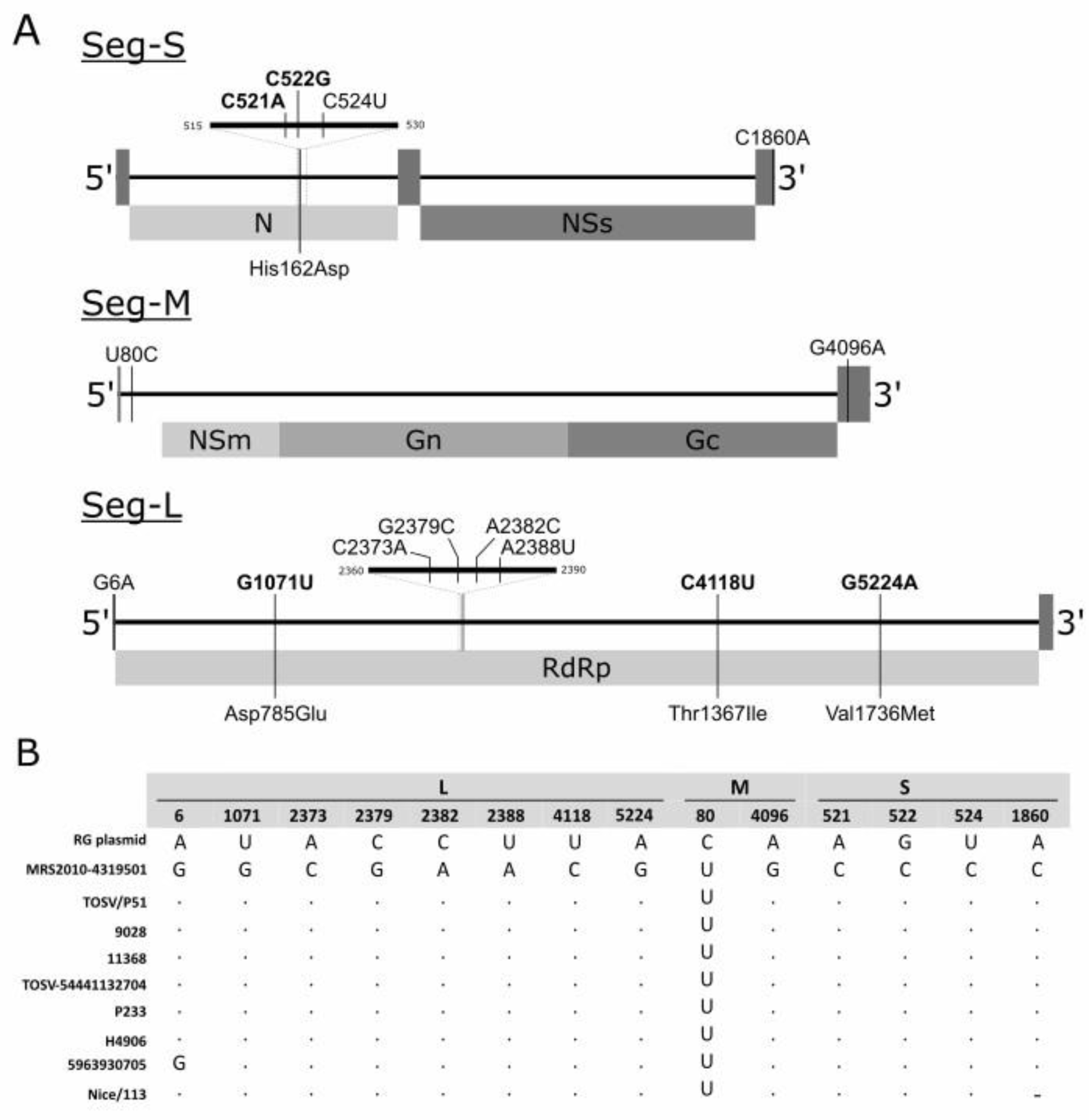
Comparison between the sequences of TOSV-B reverse genetic plasmids and those of TOSV lineage B reference genomes. (A) Schematic representation of the TOSV antigenome segments: Seg-S, Seg-M, and Seg-L. The location of the viral 5’ and 3’ untranslated regions (UTR) and open reading frame of the structural (N, Gn, Gc and RNA-dependent RNA polymerase [RdRp]) and non-structural (NSs and NSm) viral proteins are indicated. The 14 substitutions identified between the reference sequences of the wt TOSV-B and those of the plasmid developed for the reverse genetics system (indicated as RG plasmid) are presented. Non-synonymous substitutions are in bold and their respective change in amino acid is indicated. (B) Nucleotide differences between TOSV-B strains. Sequences were aligned using MUSCLE software. Dots indicate identical nucleotides. Note that, for Nice/113 strain, Seg-S sequence is incomplete at the 3’ UTR, thereby preventing comparison at position 1860.

### rTOSV-A replicates more than rTOSV-B

The replication rates of TOSV-A and TOSV-B rescued viruses (hereafter referred to as rTOSV-A and rTOSV-B) were compared to those of their parental virus strains (i.e., wt TOSV-A and wt TOSV-B) in A549 and A549 Npro cells. A549 cells constitutively express the N-terminal protease of bovine viral diarrhea virus, which blocks type I interferon (IFN-I) synthesis [16] thus allowing to assess the impact of IFN-I responses on TOSV replication. The infectious virus production by rTOSV-A, rTOSV-B, wt TOSV-A, and wt TOSV-B was determined by TCID50 assays from the supernatants of infected cells (Fig 2). Unlike rTOSV-B, rTOSV-A had higher viral titers than wt TOSV-A in both A549 and A549 Npro cells (Fig 2). Importantly, both rTOSV-B and wt TOSV-B produced lower viral titers than rTOSV-A and wt TOSV-A in both cell lines. Overall, these data indicate that rescued viruses behave similarly to their respective parental virus strains, and suggest that the difference observed between TOSV-A and TOSV-B does not depend on the IFN-I pathway.

**Fig 2.**
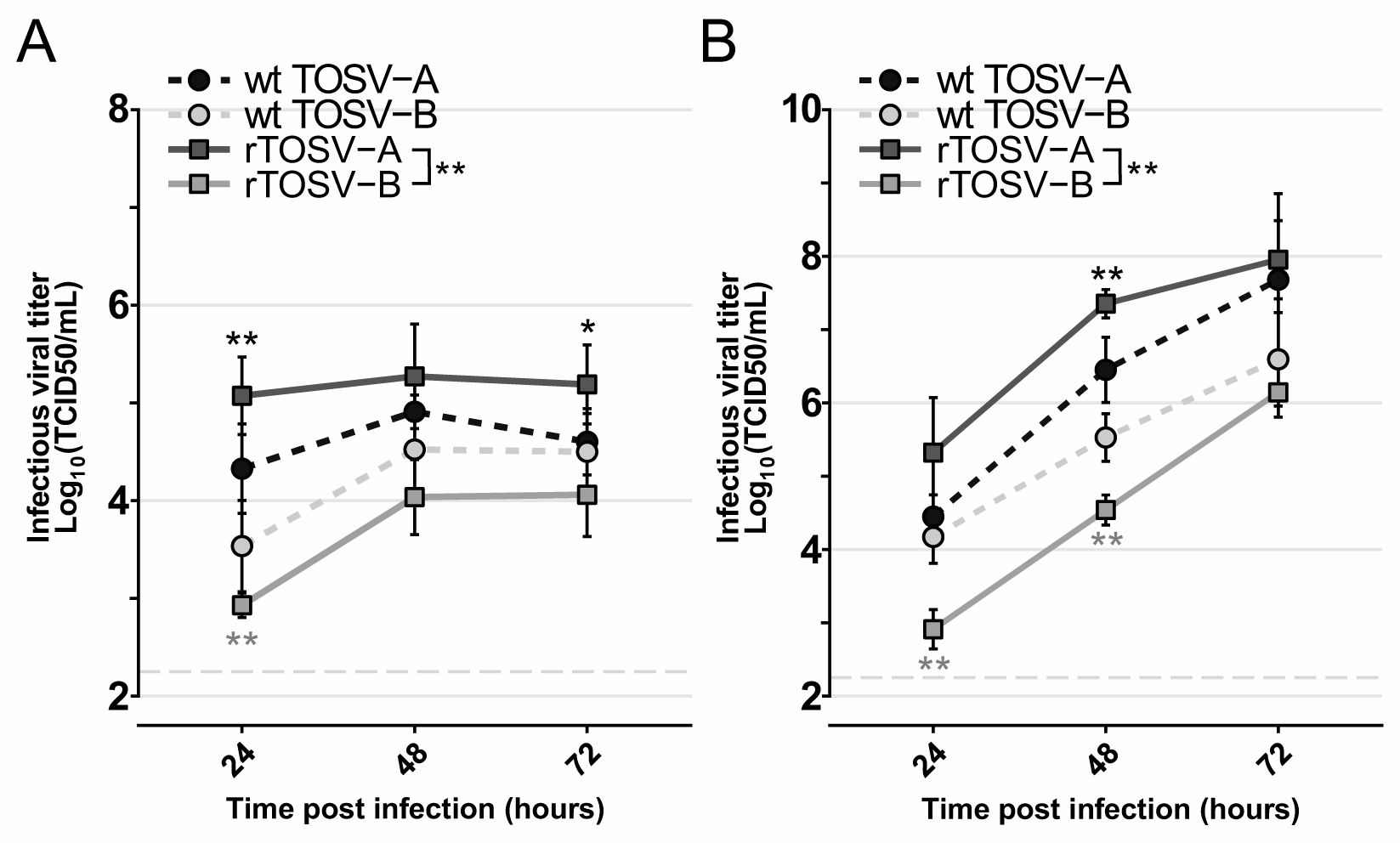
TOSV-A viruses produce higher infectious titers than TOSV-B. Growth curves of wt TOSV-A, wt TOSV-B, rTOSV-A and rTOSV-B in A549 (A) or A549 Npro (B) cells (MOI=0.01). Cell supernatants were collected at 24-, 48-, and 72-hours post-infection, and viral titers were obtained by TCID50 limiting-dilution assays in A549V cells. Each experiment was performed in triplicate and at least three times independently. Bars indicate standard deviations. Light grey dashed lines indicate the threshold of virus detection [2.25 Log_10_(TCID50/mL)]. Statistical significance was determined using multiple Wilcoxon tests with the Benjamini-Yekutieli (BY) p-value adjustment method: p<0.05 (*) and p<0.01 (**). Black and grey stars indicate the statistical significance of the rescued viruses compared to their parental virus strain (i.e., wt TOSV-A or wt TOSV-B, respectively).

### Seg-M modulates TOSV replication *in vitro*

The comparative genomic analyses of rTOSV-A and rTOSV-B revealed a high genetic diversity across the TOSV genome. In order to assess which of the three TOSV segments is responsible for the viral titer differences observed, a library of reassortant viruses was generated for each of the three genome segments, and their respective replication rates were assessed in A549 and A549 Npro cells (Fig 3A and 3B). Seg-L and Seg-S reassortant viruses had similar viral titers to those of their corresponding parental viruses (i.e., rTOSV-A or rTOSV-B) in both cell lines. However, rTOSV-A-MB (a reassortant virus carrying the Seg-S and Seg-L of TOSV-A and the Seg-M of TOSV-B) produced significantly lower viral titers than rTOSV-A (p-value <0.05 in A549 cells; p-value <0.01 in A549 Npro cells). Conversely, rTOSV-B-MA (a reassortant virus carrying the Seg-S and Seg-L of TOSV-B and the Seg-M of TOSV-A) had significantly higher viral titers than rTOSV-B (p-value <0.05 in A549 cells; p-value <0.01 in A549 Npro cells). These data demonstrate that Seg-M carries the viral determinant responsible for the higher replication rate of rTOSV-A compared to rTOSV-B.

**Fig 3.**
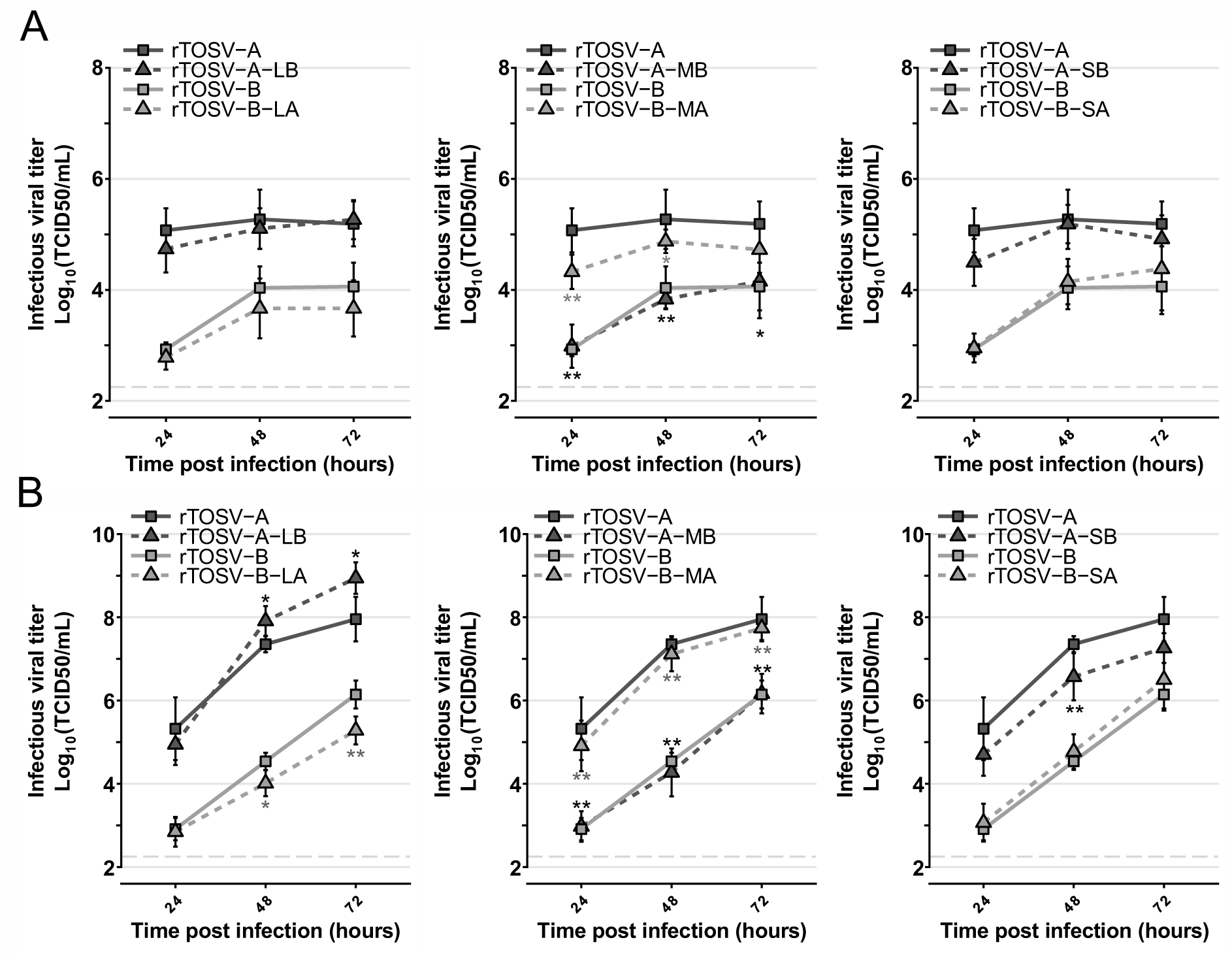
Seg-M modulates TOSV replication rates in A549 and A549 Npro cells. Growth curves of rTOSV-A and rTOSV-B and their respective Seg-L (LB or LA), Seg-M (MB or MA) and Seg-S (SB or SA) reassortant viruses in A549 (A) or A549 Npro (B) cells (MOI=0.01). Cell supernatants were collected at 24-, 48-, and 72-hours post-infection, and viral titers were determined by TCID50 limiting-dilution assays in A549V cells. Each experiment was performed in triplicate and at least three times independently. Bars indicate standard deviations. Light grey dashed lines indicate the threshold of virus detection [2.25 Log_10_(TCID50/mL)]. Statistical significance was determined using multiple Wilcoxon tests with the BY p-value adjustment method: p<0.05 (*) and p<0.01 (**). Black and grey stars indicate the statistical significance of reassortant viruses compared to their corresponding parental virus (i.e., rTOSV-A or rTOSV-B, respectively).

### The Gn-Gc coding sequence modulates TOSV replication

TOSV Seg-M encodes a polyprotein which is further cleaved by the host proteases into NSm, Gn, and Gc [17]. Genomic comparison between rTOSV-A and rTOSV-B showed a high genetic diversity distributed throughout the entire M polyprotein coding sequence (Fig 4A). Interestingly, the NSm harbours a higher genetic diversity (81% amino acid homology) compared to the Gn or Gc (89% and 95%, respectively). In order to understand which of these proteins is involved in modulating TOSV replication rate, chimeric Seg-M were produced by exchanging NSm or Gn-Gc coding sequence between rTOSV-A and rTOSV-B (Fig 4A). The replication rates of these rescued viruses were assessed in A549 Npro cells (Fig 4B-D). No significant difference was observed in the replication kinetics of the two NSm chimeric viruses (Fig 4B). However, rTOSV-A-GnGcB displayed significantly lower viral titers than rTOSV-A (p-value <0.01; Fig 4C). Conversely, rTOSV-B-GnGcA showed significantly higher viral titers than rTOSV-B (p-value <0.002; Fig 4D). Altogether, these data demonstrated that the Gn-Gc coding sequence is responsible for the higher replication rate of rTOSV-A compared to rTOSV-B.

**Fig 4.**
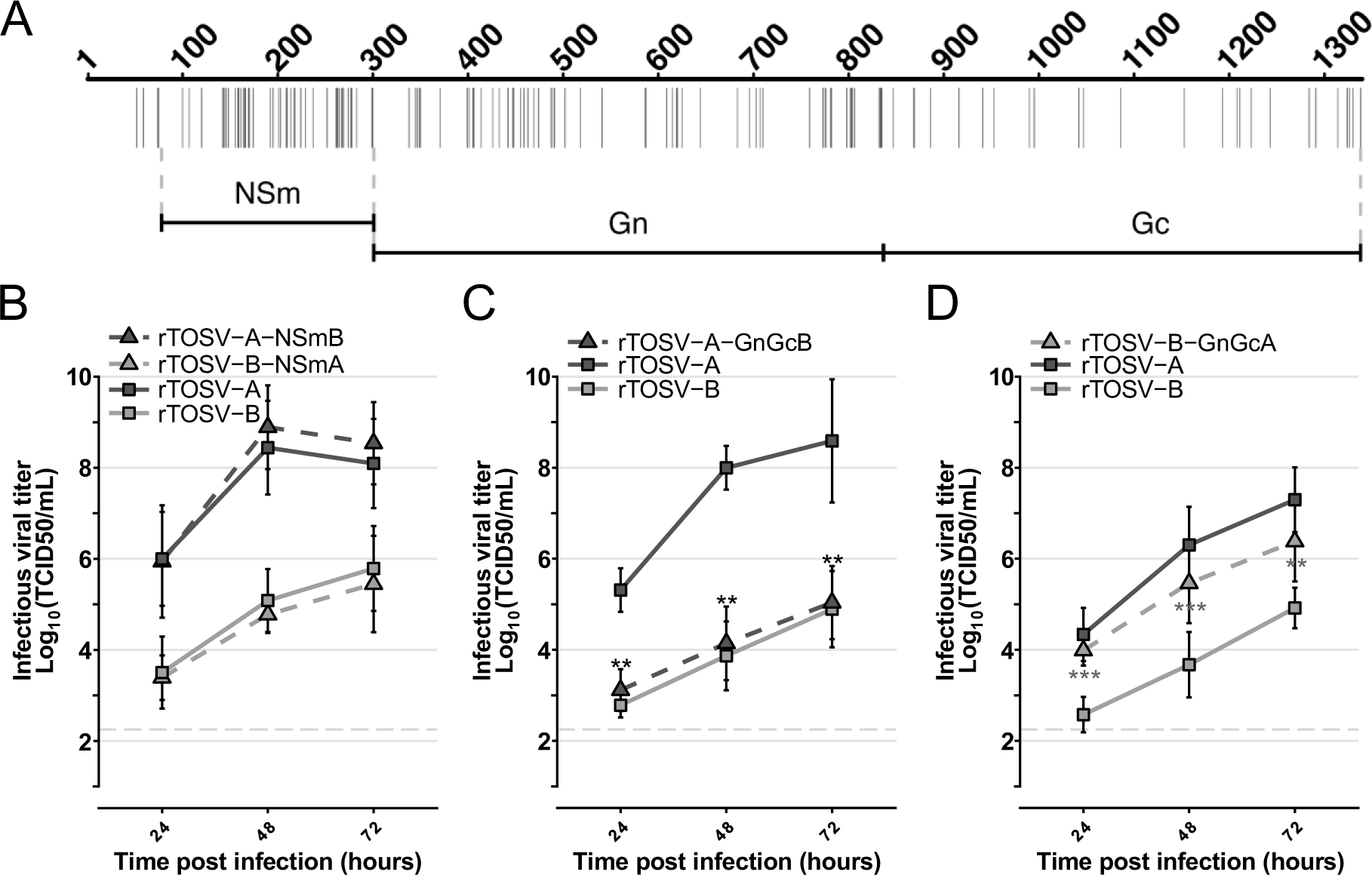
Gn-Gc modulates TOSV replication rates in A549 Npro cells. (A) Schematic representation of the M polyprotein sequence alignment between rTOSV-A and rTOSV-B. Each vertical bar represents an amino acid substitution. The region corresponding to either NSm or Gn-Gc protein is indicated. The dashed lines show the limits of the exchanged coding sequence in the NSm or Gn-Gc chimeric viruses. (B-D) Growth curves in A549 Npro cells infected with rTOSV-A, rTOSV-B or NSm (B), rTOSV-A-GnGcB (C), or rTOSV-B-GnGcA (D) chimeric viruses (MOI=0.01). Cell supernatants were collected at 24-, 48-, and 72-hours post-infection, and viral titers were obtained by TCID50 limiting-dilution assays in A549V cells. Each experiment was performed in triplicate and at least three times independently. Bars indicate standard deviations. Light grey dashed lines indicate the threshold of virus detection [2.25 Log_10_(TCID50/mL)]. Statistical significance was determined using multiple Wilcoxon tests with the BY p-value adjustment method: p<0.05 (*) and p<0.01 (**). Black and grey stars indicate the statistical significance of chimeric viruses compared to their corresponding parental virus (i.e., rTOSV-A or rTOSV-B, respectively).

### rTOSV-A virions are internalised faster than rTOSV-B in infected cells

Phlebovirus glycoproteins play a key role in virus entry [18]. In order to understand whether the genetic diversity within the Gn-Gc coding sequence of rTOSV-A and rTOSV-B could affect the viral entry process, a virus penetration kinetic assay was carried out as described in Fig 5A. Data were analysed using a 3-parameter logistic model described elsewhere [19]. Cell internalization of rTOSV-A started within the first 10 minutes following temperature shift, and it increased over time to reach the half-maximal level (t1/2) within 22.84±1.19 minutes, and a plateau 20 minutes later (Fig 5B). In contrast, rTOSV-B reached the t1/2 in 34.88±1.34 minutes (Fig 5B). These data demonstrate that the cell penetration kinetic of rTOSV-A occurs faster than that of rTOSV-B.

**Fig 5.**
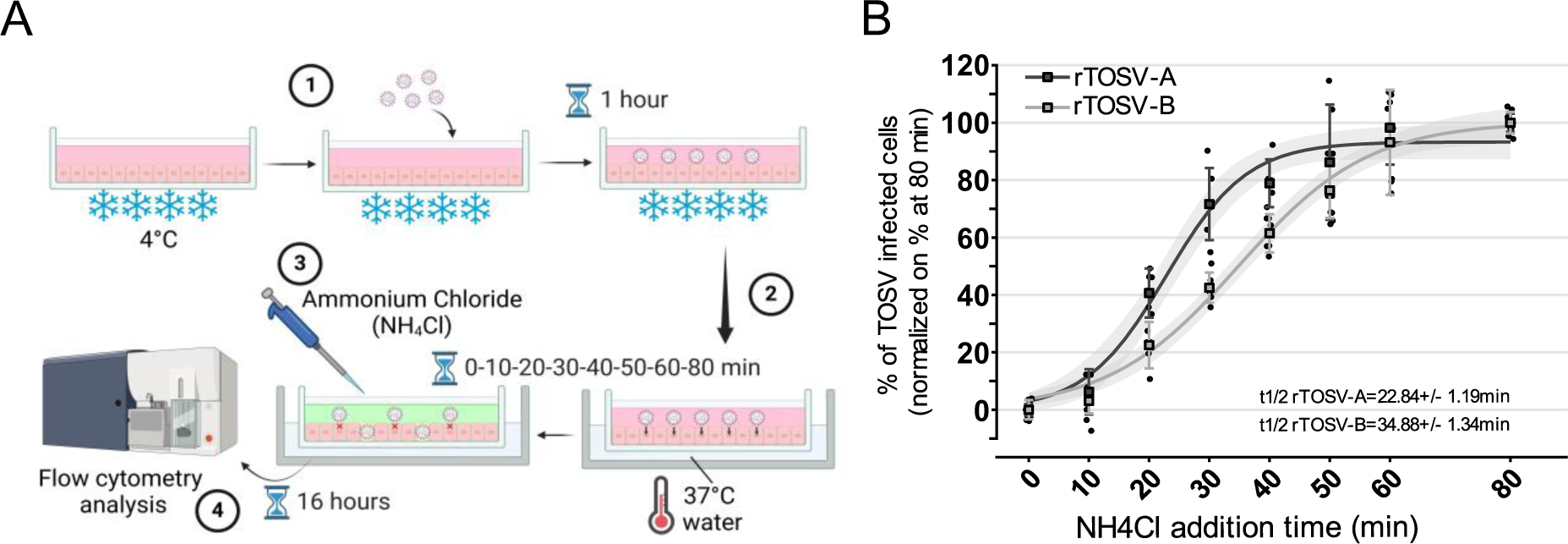
rTOSV-A virions are internalized faster than rTOSV-B virions. (A) Schematic representation of the penetration kinetic workflow used in this study. rTOSV-A and rTOSV-B virions were bound on A549 Npro cells at MOI=1 (1). Next, the temperature was rapidly increased to 37°C (2) and, at the indicated time (ranging from 0 to 80 minutes), cell culture supernatants were replaced by media containing NH_4_Cl to increase the pH and block viral fusion (3). Cells were collected at 16 hours post-infection and subsequently analysed by immunostaining (with antisera raised against TOSV) and flow cytometry (4). (B) The graphic shows the percentages of TOSV-infected cells normalized to those incubated at 37°C throughout the experiment (i.e., for 80 minutes). Penetration kinetics were modelled using a 3-parameter logistic model described elsewhere [19] and are represented in the graph as curves. The grey zones indicate the 95% confidence interval. Each experiment was performed in duplicate and at least three times independently. Bars indicate standard deviations.

### rTOSV-B Gn-Gc are more abundant than rTOSV-A Gn-Gc in infected cell supernatants

The rTOSV-A and rTOSV-B Gn and Gc expression and properties were further characterized by western blot. As shown in Fig 6A, rTOSV-B Gc molecular weight is approximately 0.9 kDa lower than that of rTOSV-A. To investigate if this difference was due to the different N-glycosylation patterns of rTOSV-A and rTOSV-B Gc proteins, rTOSV-A and rTOSV-B virions from A549 Npro infected cells were treated with PNGase-F. No difference in terms of glycosylation between rTOSV-A and rTOSV-B Gc proteins was observed (Fig S1). In addition, rTOSV-A and rTOSV-B infected cells had similar Gn and N protein levels (Fig 6A and 6B). However, rTOSV-B infected cells had significantly higher Gc levels compared to rTOSV-A infected cells (p-value <0.0001; Fig 6B). Moreover, higher Gn and Gc levels were found in the supernatants of rTOSV-B infected cells compared to rTOSV-A infected cells (Fig 6A and 6B), suggesting a higher incorporation of Gn and Gc into newly produced rTOSV-B virions.

**Fig 6.**
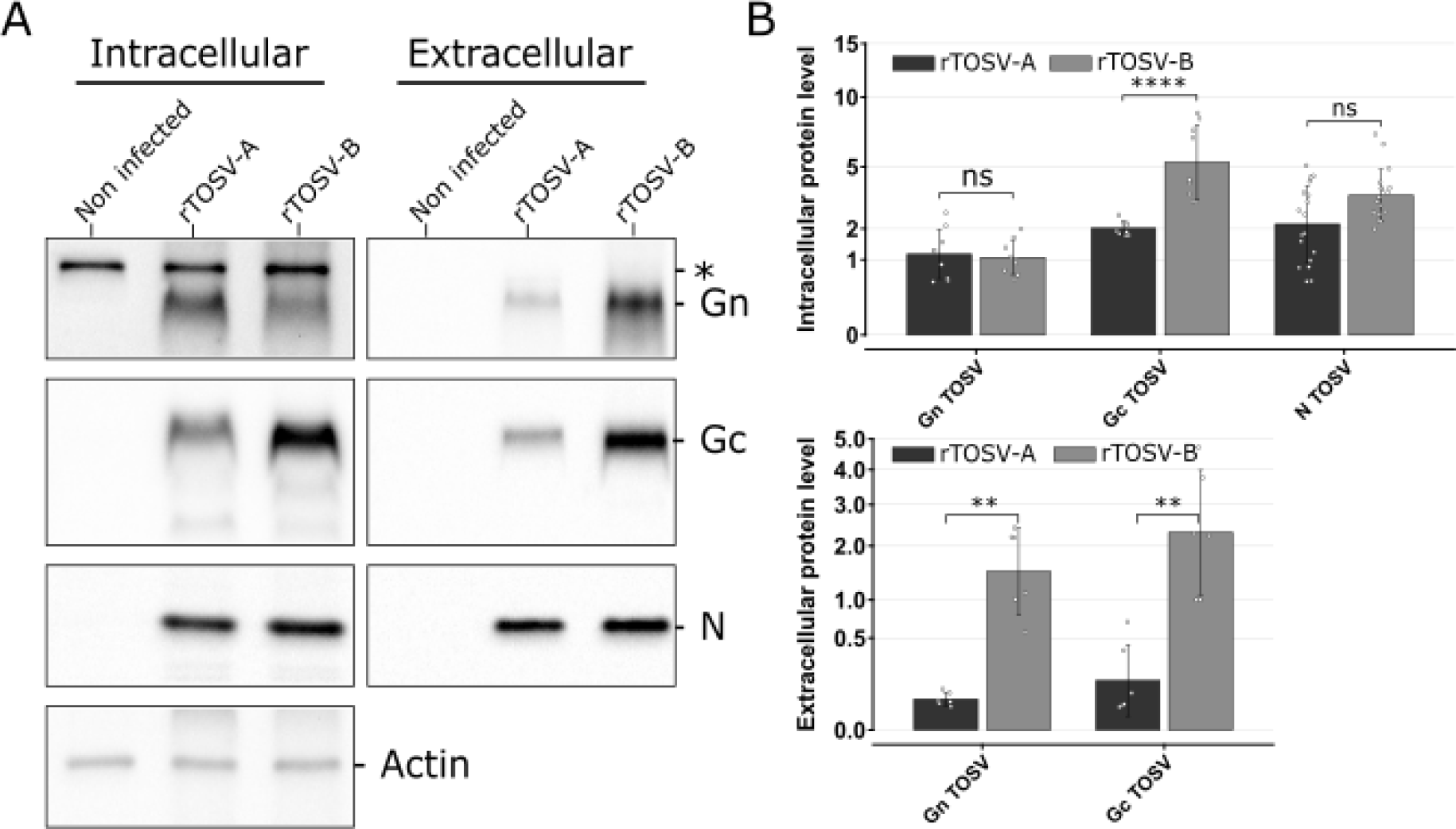
Intracellular and extracellular protein levels of rTOSV-A and rTOSV-B Gn and Gc glycoproteins. (A) Following infection of A549 Npro cells (MOI=0.1) with rTOSV-A or rTOSV-B viruses, cells and supernatants were collected at 24 hours post-infection and analysed by western blot with antisera raised against TOSV N, Gn or Gc proteins, or actin, as indicated. Each experiment was performed in triplicate and at least three times independently. Representative blots are presented. A non-specific band in cell lysates was observed with TOSV Gn antibody and is indicated by an asterisk. (B) Signals from rTOSV-A and rTOSV-B N, Gn, Gc, and actin were quantified from three independent experiments using Image Lab software. Protein levels were normalized against actin levels for the intracellular compartment or against N protein level for the extracellular fraction. Bars indicate standard deviations. Statistical significance was determined using multiple Wilcoxon tests with the BY p-value adjustment method: p>0.05 (ns: not significant), p<0.01 (**), and p<0.0001 (****).

The intracellular localisation of rTOSV-A and rTOSV-B Gn and Gc proteins was also investigated in A549 Npro cells. Both proteins displayed a perinuclear localisation, potentially reflecting localisation in the endoplasmic reticulum and/or Golgi apparatus (Fig S2).

### Newly produced rTOSV-A virions display a higher infectivity than rTOSV-B

In order to understand if the increased amount of Gn and Gc proteins incorporated into newly produced rTOSV-B virions could affect viral infectivity, physical and infectious titers of both rTOSV-A and rTOSV-B were determined in the supernatants of infected cells before the appearance of cytopathic effects. To this end, the number of rTOSV-A and rTOSV-B genomic copies was quantified by RT-qPCR, and the number of infectious viral particles was determined by virus titration assays (Fig 7). As shown in Fig 7A, rTOSV-B infected cells released more Seg-S and Seg-M than rTOSV-A, even though rTOSV-A produced more infectious viral particles than rTOSV-B (Fig 7B). To assess the infectivity of the physical viral particles produced by rTOSV-A and rTOSV-B, the ratio of genomic copies *versus* the number of infectious viral particles was calculated (Fig 7C). Our results show that, regardless of the viral segment considered, this ratio is significantly higher for rTOSV-B than rTOSV-A (p-value <0.01; Fig 7C), suggesting that rTOSV-A is more efficient at producing infectious virus particles than rTOSV-B.

**Fig 7.**
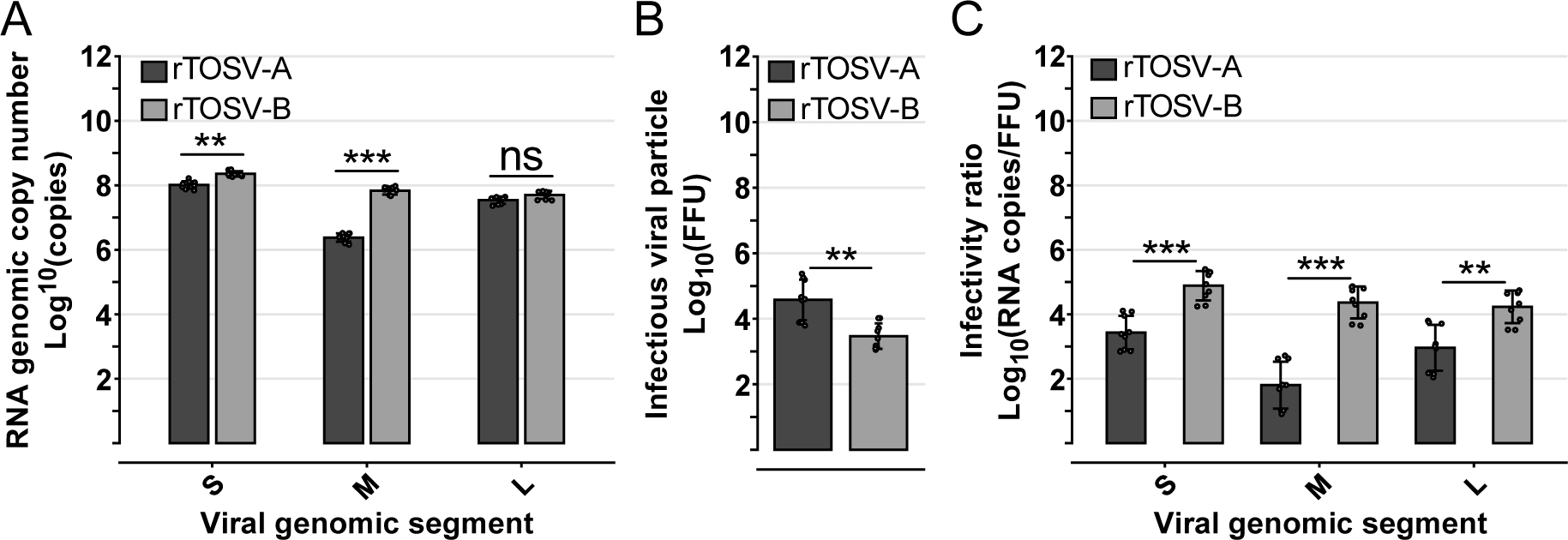
rTOSV-A virions are more infectious than rTOSV-B. A549 Npro cells were infected with rTOSV-A or rTOSV-B (MOI=0.2). Cell culture supernatants were collected at 24 hours post-infection, before the appearance of cytopathic effects. (A) RNA copy numbers of Seg-S, Seg-M, and Seg-L of rTOSV-A and rTOSV-B were quantified by RT-qPCR. (B) Infectious viral particle number was determined by foci forming assay and expressed as Log_10_(FFU). (C) The infectivity of viral particles was estimated by calculating the ratio of genomic RNA copies against infectious viral titers. Each experiment was performed in triplicate and at least three times independently. Bars indicate standard deviations. Statistical significance was determined using multiple Wilcoxon tests with BY p-value adjustment method: p>0.05 (ns: not significant), p<0.01 (**), and p<0.001 (***).

### rTOSV-A and rTOSV-B virions differ in size and structure

Finally, the structure of rTOSV-A and rTOSV-B virions was analyzed by transmission electron microscopy. As previously shown for other bunyaviruses [20], rTOSV-A and rTOSV-B virions were detected within the Golgi apparatus of TOSV-infected cells (Fig 8A and 8B). Intriguingly, some differences in the cell supernatants were observed at the surfaces of rTOSV-A and rTOSV-B virions (Fig 8C). More specifically, the surfaces of rTOSV-B virions appeared to be more structured and organised than those of rTOSV-A virions (Fig 8C). Furthermore, while no significant difference was observed in the size of intracellular virions, a significant difference of 20 nm was measured between rTOSV-A and rTOSV-B virions in the supernatants of infected cells (p-value <0.0001; Fig 8D).

**Fig 8.**
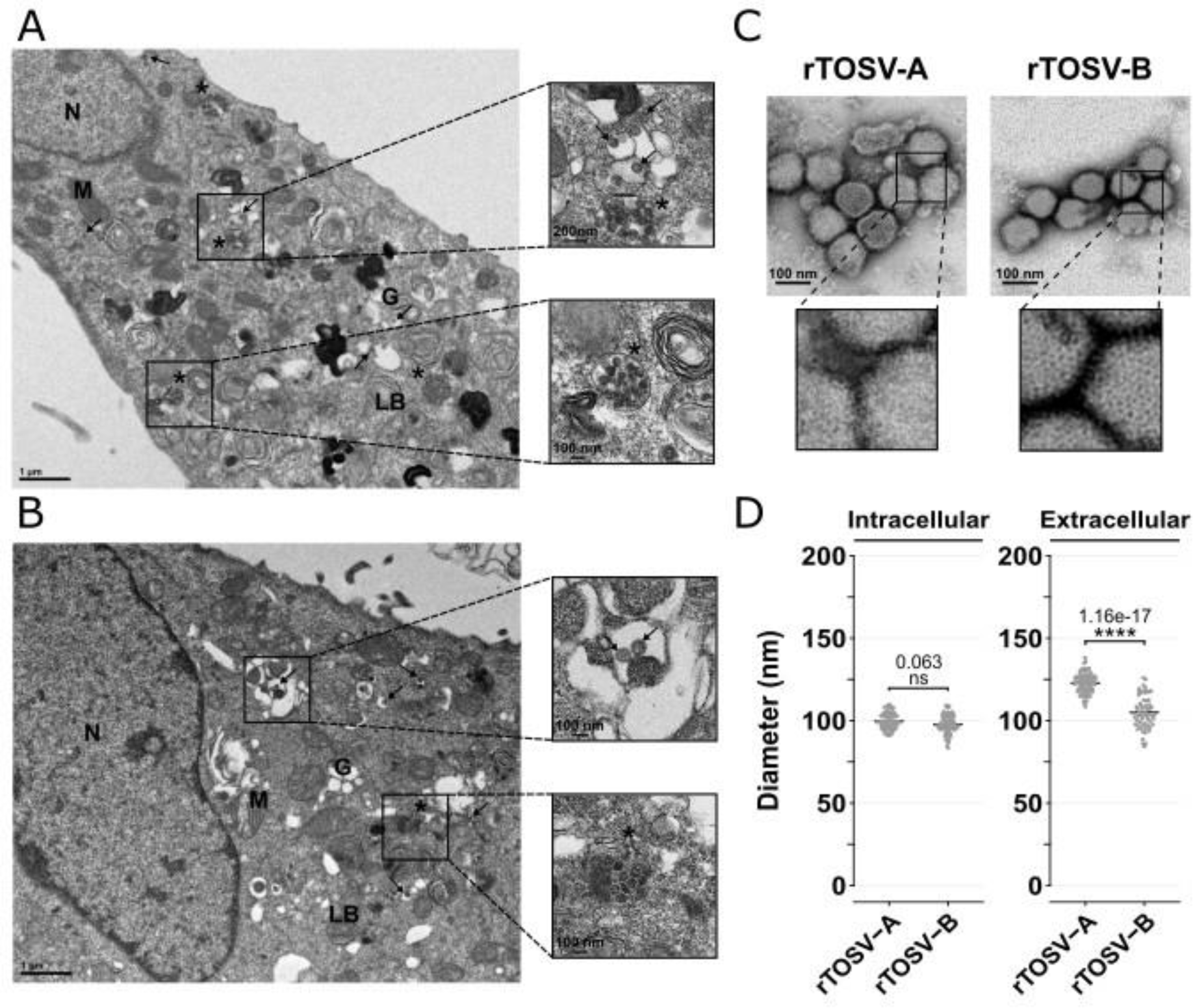
rTOSV-A and rTOSV-B virions differs in surface structure and size. (A-B) A549 Npro cells were infected with rTOSV-A (A) and rTOSV-B (B) viruses (MOI=0.1). Cells were fixed at 24 hours post-infection, before the appearance of cytopathic effects, and prepared for ultrastructural study as described in Materials and Methods. Ultrathin sections of cell pellets were cut, collected on grids, and stained before observations under transmission electron microscope. Representative pictures of rTOSV-A or rTOSV-B infected cells are shown. The experiment was performed in duplicate; at least 15 different infected cells were imaged per condition. Arrows indicate virions present within the Golgi apparatus. Unidentified vesicles containing electron-dense virus-like structures are indicated with an asterisk. N, nucleus; M, mitochondria; G, Golgi apparatus; LB, lamellar body. (C) rTOSV-A and rTOSV-B virions from cell supernatants were purified, and the viral particles were observed by transmission electron microscopy after negative staining. Representative pictures of rTOSV-A or rTOSV-B virions are shown. (D) The diameter of viral particles detected inside the cells or in the cell supernatants was determined. At least 60 different virions were analysed per condition. Statistical significance was determined using T-test with BY p-value adjustment method: p>0.05 (ns: not significant) and p<0.0001 (****).

## Discussion

In this study, we developed a new reverse genetic system for wt TOSV-B to carry out a comparative approach between two genetically distinct TOSV strains. This allowed us to unveil that the genetic diversity of TOSV significantly impacts its biological properties. More specifically, we demonstrated that two TOSV strains belonging to two lineages A and B displayed different replication kinetics in mammalian cell lines. We showed that TOSV-A and TOSV-B have different Gn and Gc coding sequences, and that these sequences contain the determinant(s) responsible for the different viral replication kinetics.

Gn and Gc proteins are key viral proteins involved in phlebovirus entry [18]. Along these lines, our results showed that rTOSV-A penetrates cells faster than rTOSV-B. Further analyses on rTOSV-A and rTOSV-B entry, similar to those already performed elsewhere [21], would help us defining more precisely at which step of this process these two TOSV viruses differ.

Importantly, the infectivity of newly produced viral particles was found to be higher for rTOSV-A than rTOSV-B. Due to the segmented nature of bunyaviruses genomes, the generation of infectious progeny relies on an efficient genome packaging that is thought to be modulated by the Gn protein [22,23]. Interestingly, although rTOSV-B displayed higher levels of Gn-Gc proteins and genomic RNA copies than rTOSV-A, it produced lower levels of infectious virus particles. These differences may be due to a less efficient genome packaging process in rTOSV-B than rTOSV-A due to the amino acid differences found in the glycoprotein coding sequences of these two viruses. Further experiments are needed to identify which amino acid(s) of the Gn or Gc protein is (are) involved in the differences observed, and to what extent these differences are specific to these two strains or, more generally, to the two genetic lineages.

Interestingly, we observed a difference in the size of Gc and in the amount of Gn and Gc in the supernatants of cells infected with rTOSV-A or rTOSV-B. The reason for the different molecular weight of Gc proteins is not yet known, though it would appear unrelated to N-glycosylation. One can speculate that these two proteins are cleaved differently: a longer rTOSV-A Gc or a shorter rTOSV-B Gc protein could in turn affect their folding and interaction with Gn, resulting, respectively, in reduced or increased glycoprotein incorporation on the surface of viral particles, and/or in morphological changes of the virions. Additionally, a lower level of Gn-Gc incorporation in rTOSV-A particles compared to rTOSV-B may result in a more relaxed virion structure, explaining the larger size and rougher/more irregular surface of rTOSV-A compared to rTOSV-B virions. Lastly, the amount of viral glycoproteins incorporated at the surface of virus particles can affect virus entry, as shown for RVFV, where higher incorporation of the p78 glycoprotein results in reduced replicative capacity and lower virus binding to the cell surface, ultimately leading to inefficient virus internalisation [24]. Thus far, no high-resolution structure of TOSV particles has yet been described, making it impossible to predict how much the sequence diversity between Gn and Gc of TOSV-A and TOSV-B could affect virion structure. Further investigations using high-resolution cryo-electron microscopy would be required to analyse these differences.

Although no inter-lineage reassortant viruses have yet been reported for TOSV, their existence was described for numerous species belonging to the *Bunyavirales* [25]. Notably, in this study, we showed that all segment combinations resulted in efficient virus rescue *in vitro*. It therefore appears that reassortment between TOSV-A and TOSV-B is possible in nature, especially given that these two TOSV strains and lineages are co-circulating in France [14,15]. Importantly, the vast majority of reassortments described to date involve the exchange of the Seg-M [26], and we identified this segment as a key element in modulating infection by the two TOSV strains studied. Hence, TOSV reassortment, especially those involving Seg-M, should be monitored more closely as such an event may significantly alter the biology of the virus in terms of pathogenicity. The characterisation of circulating TOSV strains by sequencing is therefore important and represents a challenge for the surveillance of this virus. Comparative approaches between different strains of TOSV, such as those carried out in this study, are thus crucial to determine to what extent and how the genetic diversity of TOSV influences its biology. Moreover, the data gathered from this kind of approaches may be relevant to better characterize the pathogenicity of the disease and develop vaccine strategies taking into account TOSV glycoproteins genetic diversity.

## Materials and Methods

### Cells

BSR, BSR T7/5 CL21, VeroE6, A549 and A549 Npro and A549V cells were grown in Dulbecco’s Modified Eagle Medium (DMEM) (Gibco, Thermo Fisher Scientific, France), supplemented with 10% heat-inactivated foetal bovine serum (FBS) (GE HEALTHCARE Europe GmbH, Germany). All cell lines were cultured in a 37°C, 5% CO_2_ humidified incubator. BSR cells are derived from the BHK-21 cells and were kindly provided by Karl-Klaus Conzelmann (Ludwig-Maximilians-University Munich, Gene Center, Munich/Germany). BSR T7/5 CL21 cells are a clone from the BSR T7/5 cell line [27]. VeroE6 cells were obtained from Philippe Marianneau (Unité de virologie - ANSES Lyon, France). A549 Npro and A549V cells were obtained from Richard E. Randall (University of St. Andrews, UK). A549 Npro cells are derived from the A549 cell line and are constitutively expressing the N protein of the bovine viral diarrhea virus that blocks the interferon synthesis pathway [16]. In A549V cells, stimulation by IFN-I is inhibited by the expression of the V protein of the simian virus 5 [28].

### Production of rTOSV-B reverse genetic plasmids

Viral RNA was extracted from TOSV MRS2010-4319501 (wt TOSV-B) [15] (obtained from EVAg, France) BSR infected cells using Trizol reagent (Thermo Fisher Scientific, France) following the manufacturer’s protocol. RNA was then purified using the RNeasy Plus Mini kit (QIAGEN, Germany). Complementary DNA (cDNA) was synthesized using Omniscript RT polymerase (QIAGEN, Germany) and viral genomic segment-specific primers. Primers and viral RNAs were first incubated for 10 minutes at 65°C before the RT reaction following manufacturer protocol. Full-length segments were then amplified from cDNA by PCR using CloneAmp HIFI PCR Premix (Takara, France) and then gel purified using NucleoSpin Gel and PCR Clean-up kit (Macherey-Nagel, Germany). The plasmid backbone, containing the T7 promoter and terminator, the HDV ribozyme and the ampicillin resistance gene, was amplified by PCR from the pUC57 plasmid used in the TOSV reverse genetic system described previously [12]. Linearized plasmids were digested for 1 hour at 37°C by DnpI (New England Biolabs, UK) prior to performing the cloning reaction using the In-Fusion HD-Cloning Plus kit (Takara, France) according to the manufacturer’s instruction with a molar ratio insert:vector of 2:1 for the S and M segment and a 1:2 molar ratio for the L segment. Stellar competent bacteria (Takara, France) were used for transformation and plasmid amplification following manufacturer protocol. To construct chimeric plasmids, the NSm or Gn-Gc coding sequences were amplified by PCR from the reverse genetic plasmid of TOSV-A or -B, using primers with 15 bases homologous to the end of the target plasmids (either pUC57-TOSV_M-ΔNSm or -ΔGn-Gc) at their 5’ end. Additionally, pUC57-TOSV_M-ΔNSm or -ΔGn-Gc plasmids were generated by PCR on plasmids pUC57-TOSV-A_M or pUC57-TOSV-B_M. Chimeric plasmids were then ligated and amplified using In-Fusion cloning process as described above. The sequences of all generated plasmids were verified by sequencing. All RT and PCR primer sequences are available on request.

### Validation of the plasmid’s sequences

After sequencing, plasmid sequences were compared to wt TOSV-B strain and other lineage B TOSV strains (TOSV/P51: KU204978-KU204980, TOSV 9028: KU904263-KU904265, TOSV 11368: KU925897-KU925899, TOSV 54441132704: KX010932-KX010934, TOSV P233: KU204975-KU204977, TOSV H4906: KU922125-KU922127, TOSV 5963930705: KU935733-KU935735, and TOSV Nice/113: KU204981-KU204983) using MUSCLE software [29]. For each nucleotide substitution in relation to the deposited sequence of wt TOSV-B, RNA was extracted from the viral stock, the region concerned was amplified, and then sequenced.

### Rapid amplification of cDNA-ends by polymerase chain reaction (RACE-PCR)

The genome and antigenome 3’ termini were sequenced using RACE-PCR analysis as described before [12]. Briefly, RNA was extracted from viral stock using QIAmp viral RNA (QIAGEN, Germany), polyadenylated with *E. coli* Poly(A) Polymerase I from mMESSAGE mMACHINE™ T7 ULTRA kit (Invitrogen, by Life Technologies, Thermo Fisher Scientific, France) before being purified using RNeasy® Plus Mini kit (QIAGEN, Germany). Complementary DNA was synthesized using the PrimeScript RT reagent kit (Takara, France) and oligo-d(T)-AP primer (GACCACGCGTATCGATGTCGACTTTTTTTTTTv). Seg-S termini were then amplified using anchor primer (GACCACGCGTATCGATGTCGAC) and RACE_TOSV_S1 (CCACCTCATTAGCCTGCTTAG) or RACE_TOSV_S2 (GTCACTCTCTTGTCTTCCTTAG). Seg-M termini were amplified using anchor primer and RACE_TOSV_M1 (TTCCTCTTCCGTTCCAGCTC) or RACE_TOSV_M2 (CCATATTCTCCAGAGCCTCG). Seg-L termini were amplified using anchor primer and RACE_TOSV_L1 (TCTCCAGTCTGCTGACAGTC) or RACE_TOSV_L2 (TCCTGACTCAGAGCCTGAAG). Sequences were next determined using Sanger sequencing. Site-directed mutagenesis was performed on the 3’ antigenomic terminus of the Seg-S and the 5’ antigenomic terminus of Seg-L in order to align plasmid sequences with those obtained from the RACE-PCR analysis. Briefly, after a 12-cycle PCR, PCR products were treated with DpnI and purified on agarose gel. Plasmids were then transformed and amplified as described above, and the plasmid sequences were reassessed by Sanger sequencing. Complete sequences of each segment used for the reverse genetics system (Seg-L, Seg-M and Seg-S) are accessible under the GenBank accession numbers PQ015116, PQ015117 and PQ015118, respectively.

### Virus production by reverse genetics

Viral rescue was carried out by transfection using Lipofectamine 2000 (Invitrogen by Life Technologies, Thermo Fisher Scientific, France). BSR T7/5 CL21 cells were seeded in six-well plates at a density of 3×10^5^ cells per well and incubated for 24 hours to reach 80% of confluency. Culture media was changed to 1mL of DMEM complemented with 4% of FBS. Cells were then transfected with 500ng of each plasmid expressing the S, M and L RNA antigenome under the control of a T7 RNA polymerase promoter by using 3µL of Lipofectamine 2000 in a final volume of 60µL of OptiMEM (Gibco, Thermo Fisher Scientific, France). After incubation for 6 hours at 37°C and 5% CO2, cells were washed with 1mL of phosphate-buffered saline (PBS; Sigma-Aldrich, Merck, France). Then, 2ml of DMEM supplemented with 4% FBS was added to each well, and the cells were further incubated at 37°C until cytopathic effects appeared. The cell supernatants, referred to as passage 0 (p0), were then collected, clarified by centrifugation and stored at −80°C.

### Viral stock production

All viral stocks were amplified from p0 on VeroE6 cells for 2 to 3 days in DMEM supplemented with 4% FBS at 37°C and 5% CO_2_. For high MOI infection assays, viral stocks were concentrated by ultracentrifugation. Briefly, VeroE6 were seeded 3 days prior to infection using p0. Four days later, media was collected and clarified by centrifugation before ultracentrifugation on a 20% (g/mL) ultrapure sucrose diluted in PBS (Invitrogen by Life Technologies, Thermo Fisher Scientific, France). Ultracentrifugation was performed in an Optima XE-90 machine (Beckman Coulter Life Sciences, USA) at 124000g for 3 hours at 4°C. Pellets were then resuspended in PBS and stored at −80°C.

### Foci forming titration assays

Viral stock titers were determined by foci-forming assays using VeroE6 cells. Briefly, VeroE6 cells seeded in 12-well plates were infected with 10-fold serial dilutions of virus stocks and incubated for 2 hours at 37°C. The media was then removed and wash twice in PBS. Cells were then covered with 1.5mL of a semisolid overlay containing a volume-to-volume mixture of 2.5% ultrapure agarose (Life Technologies, Thermo Fisher Scientific, France) and 2X minimal essential medium (MEM; Gibco, Thermo Fisher Scientific, Villebon-sur-Yvette, France) supplemented with 4% FBS and incubated at 37°C for 6 days. Cells were then fixed in 4% formaldehyde (Formaldehyde 37-41% diluted to 4% in ultra-pure water, Fisher Chemical, Thermo Fisher Scientific, USA) overnight at 4°C, before agarose overlay removal and permeabilization with 0.1% Triton (Euromedex, France) in PBS. A first blocking step was performed with a solution consisting of PBS containing 0.1% Tween 20 (Euromedex, France) and 0.4% gelatine fish (Sigma-Aldrich, Merck, Germany), prior to a second blocking using a solution with 0.1% Tween 20 and 2.5% FBS in PBS. Cells were then incubated overnight at 4°C and under agitation with mouse ascites raised against TOSV (1:2500 in second blocking solution) kindly supplied by Philippe Marianneau (Unité de virologie, ANSES Lyon, France). Infection foci were visualised using the appropriate secondary antibody coupled with horseradish peroxidase (HRP; 1:5000, Sheep Anti-Mouse IgG A5906, Sigma-Aldrich, Merck, Germany) and TrueBlueTM chemical (SeraCare, LGC Clinical Diagnostics, USA). Infection foci were then counted to determine the viral titer expressed as FFU/mL (Foci Forming Unit).

### *In vitro* growth curves

Monolayer of A549 or A549 Npro cells were seeded 24 hours before infection in 12-well plates (2.5×10^5^ cells per well). Cells were then incubated at 37°C and 5% CO_2_ with the different viruses at a multiplicity of infection (MOI) of 0.01 in DMEM complemented with 4% FBS. After two hours of infection, cells were washed twice in PBS and further incubated at 37°C in DMEM 4% FBS. At 24-, 48- and 72-hours post-infection, 100µl of the cell supernatants were collected for viral titration by limiting dilution assays in A549V cells. Briefly, in a 96-well plate, 11µL of cell supernatants were diluted in quadruplicate by a 10-fold serial dilution (from 10^−1^ to 10^−12^) in 100µL of DMEM supplemented with 4% FBS. Then, 4×10^3^ A549V cells were added to each well and incubated for 6 days at 37°C in 5% CO_2_. Cytopathic effects were assessed under a microscope, and viral titers were expressed as 50% tissue culture infective doses (TCID50/mL) using the Reed and Muench method [30]. Each experiment was done in triplicate and repeated at least three times independently, using two independent viral stocks.

### Virus entry kinetic

A549 Npro cells were seeded 24 hours post-infection in 48-well plates (7.5×10^4^ cells per well). Cells were first incubated for 10 minutes on ice in cold DMEM supplemented with 10% FBS and 20mM HEPES buffer (Gibco, Thermo Fisher Scientific, USA). rTOSV-A or rTOSV-B infectious viral particles (ranging from 7.5×10^4^ to 15×10^4^) were bound to the cells on ice for 1 hour in cold DMEM supplemented with 0.2% FBS and 20 mM HEPES buffer to synchronise the virus attachment. Then, the cell supernatants were replaced by warm DMEM medium supplemented with 4% FBS and 20mM HEPES buffer to trigger virus internalisation. At specific times after the temperature shift, the culture medium was replaced with warm DMEM supplemented with 10% FBS, 20mM HEPES buffer and 50mM ammonium chloride (NH_4_Cl, Merck, Germany). The addition of this weak base results in a rapid increase in pH within the cell [31], which inhibits the TOSV acidification-dependent fusion process in endosomes [21]. At 16 hours post-infection, the medium was removed, and the cells were collected and fixed with a 4% paraformaldehyde solution (PFA, ChemCruz, Santa Cruz Biotechnology, Germany). The percentage of infected cells was determined by flow cytometry. To this end, fixed samples were permeabilised with a solution of PBS containing 0.1% Saponin (Sigma-Aldrich, Merck, Germany) and 10% FBS. Cells were then labelled with mouse ascites directed against TOSV (1/1000) followed by a secondary antibody directed against mouse antibodies coupled to the Alexa-488 fluorochrome (1/1000, A-11001, Invitrogen by Life Technologies, Thermo Fisher Scientific, France). Primary antibodies were incubated for 2.5 hours at 37°C. Secondary antibodies were incubated for 1 hour at 37°C. All antibody solutions were diluted in PBS 0.1% saponin 10% FBS. Cells were then analysed using the Canto II cytometer (Becton Dickinson BD, USA). Data acquisition and analysis were performed using BD FACSDivaTM software (version 6, Becton Dickinson BD, USA). Each experiment was done in duplicate and repeated three times independently, using two independent viral stocks. The virus penetration kinetics were then determined using a 3-parameter logistic model as described by Fontaine *et al.* [19]. Model parameters were estimated for each condition using a non-linear least squares method based on the Gauss-Newton algorithm. Modelling was carried out using R (version 4.2.3) and R Studio (version 2023.03.0+386).

### Western blotting

A549 Npro cells were infected at an MOI of 0.1 with rTOSV-A or rTOSV-B viruses. At 2 hours post-infection, the cells were washed successively with PBS and DMEM culture medium supplemented with 4% FBS before further incubation at 37°C and 5% CO_2_. At 24 hours post-infection, culture media was ultra-centrifugated as described above and the viral pellet was resuspended in Laemmli buffer (Alfa Aesar, Thermo Fisher Scientific, USA) diluted in PBS. Cells were then washed once in PBS and lysed using Laemmli buffer. Extracellular and intracellular cell lysates were heated at 90°C for 10 minutes and stored at −20°C. Proteins were separated by SDS-PAGE on 10% polyacrylamide gel and transferred to nitrocellulose membranes (Trans Blot Turbo RTA transfer kit midi, BioRad, Marnes-La-Coquette, France). Membranes were blocked in tris buffer saline (TBS) with 0.1% Tween and 5% dry milk (Euromedex, Souffelweyersheim, France) solution, then incubated overnight at 4°C with a polyclonal guinea pig primary antibody against TOSV Gn (1:1000), a polyclonal guinea pig antibody against TOSV Gc (1:2000), or a mouse ascites raised against TOSV (1:2500). Actin was detected with monoclonal mouse antibody coupled with horseradish peroxidase (HRP; 1:75,000; A3854, Sigma-Aldrich, Merck, Saint-Quentin Fallavier, France). Membranes were exposed to the appropriate peroxidase-conjugated secondary antibodies (1:5000 goat anti-guineapig IgG A108P or 1:10000 sheep anti-mouse IgG A5906; Sigma-Aldrich, MERCK) for 1.5h and bands were visualized by chemiluminescence using the Clarity ECL Western blotting substrate (BioRad) and ChemiDoc XRS+ System (BioRad). Band intensity was quantified using Image Lab software (Bio-Rad). Each experiment was done in triplicate and repeated three times independently, using two independent viral stocks.

### Genome quantification by RT-qPCR

A549 Npro cells were infected at an MOI of 0.2 with rTOSV-A or rTOSV-B viruses as described above for western blotting. At 24 hours post-infection, culture media were collected. Part of the cell supernatants was stored at −80°C for FFU titration assays. For TOSV genome quantification, extracellular RNAs were extracted from the cell supernatants using the QIAmp viral RNA kit (QIAGEN, Germany). Infected cells were lysed in RLT plus buffer (QIAGEN, Germany), and RNAs were extracted using the RNeasy Plus Mini kit (QIAGEN, Germany). RT reactions were performed using the PrimeScript™ RT kit (Takara, France) and the following primers: TOSV_RT_S (ACACAGAGATTCCCGTGTATTAATC) for Seg-S, TOSV_RT_M (AGTCCATCCCAAGCACAA) for Seg-M and TOSV_RT_L (CTGGCAAGGGAYATCTTTGA) for Seg-L. The previously published primers STOS-50F and STOS-138R [32] were used for qPCR on Seg-S. Primers MTOS-F (CATWGAGCAGATGAAGATGAG) and MTOS-R (AGCACTTAGAGGCTGATGATG) or LTOS-F (GATTGAGAGAGAYAGTCTCTACAC) and LTOS-R (GAACTCATAACTCATCATCTCTGC) were used for quantification of Seg-M and Seg-L respectively. The qPCR reactions were performed using the TBGreen Takara SYBR qPCR kit (Takara, France) following the manufacturer’s protocols. A viral genomic RNA standard for each rTOSV-A or rTOSV-B Seg-S, M and L was produced using the T7 RiboMAX™ kit (Promega, France) and a PCR template containing a T7 promoter amplified from each reverse genetic plasmid. DNA contaminants were removed using the TURBO DNA-free™ kit (Invitrogen by Life Technologies, Thermo Fisher Scientific, France). Each experiment was done in triplicate and repeated at least three times independently, using two independent viral stocks.

### Transmission electron microscopy

A549 Npro cells were infected in a 6-well plate at an MOI of 0.1 in duplicate, as described before. At 24 hours post-infection, infected cells were fixed with 2% glutaraldehyde (Electron Microscopy Science, USA) in 0.1 M sodium cacodylate buffer (pH 7.4) at room temperature for 30 min. After three washes in 0.2 M sodium cacodylate buffer, cells were post-fixed with 1% osmium tetroxide (Electron Microscopy Science, USA) at room temperature for 1 hour, dehydrated in a graded series of ethanol and embedded in Epon. After polymerization, ultrathin sections (100 nm) in 0.15 M sodium cacodylate (pH 7.4) buffer were cut using an ultramicrotome UCT (Leica Microsystems, France) and collected on 200 mesh grids. Sections were stained with uranyl acetate and lead citrate before observations on a 1400JEM transmission electron microscope (JEOL, Japan) equipped with an Orius camera and Digital Micrograph. For extracellular virions analysis, FBS and PBS were first ultra-centrifugated at 100000g for 18 hours before filtration (0.2μM) to prepare exo-free FBS and ultrapure PBS. A549 Npro cells were seeded in 175cm² culture flasks (1×10^7^ cells per flask) 24 hours before infection with rTOSV-A or rTOSV-B which were diluted in DMEM supplemented with 4% exo-free FBS. Four days later, the cell supernatants were collected and clarified by successive centrifugation and filtration (2000g for 20 min, 5250g for 30 min, and filtration with a 0.45μm filter followed by a further filtration with a 0.2μm filter). Clarified cell supernatant was then fixed with 2% PFA (10mL of 8% EM grade PFA, Electron Microscopy Science, USA). The fixed viral particles were then purified by ultra-centrifugation at 124000g for 2 hours, resuspended in 30mL of ultrapure PBS and ultra-centrifugated a second time as described above. The pellets were then resuspended in ultrapure PBS and adsorbed on 200 mesh nickel grids coated with formvar-C for 2 min at RT. Then, grids were coloured with 2% phosphotungstic acid for 2 min and observed with the transmission electron microscope described before. Virion diameters were measured using Fiji software [33], and their size in nm was calculated by setting the software pixel/mm ratio on the scale of each image.

### Statistical analysis

Statistical analyses were performed using R and RStudio software and the rstatix package (https://CRAN.R-project.org/package=rstatix). If data followed a normal distribution, the ANOVA test was first used with a significance threshold of 0.01. Multiple T-tests with p-value adjustment [34] were used as follow-up tests. If not, the Kruskal–Wallis test was first used with a significance threshold of 0.01. In case of statistical significance, data were then compared between conditions using multiple Wilcoxon–Mann–Whitney tests as a follow-up test with Benjamini-Yekutieli (BY) p-value adjustment. Graphical representation was realized using ggplot2 package (https://CRAN.R-project.org/package=ggplot2).

## Acknowledgements

We acknowledge the contribution of the flow cytometry platforms of SFR BioSciences Gerland Lyon Sud (UMS3444/US8) and SFR Santé Lyon-Est (UAR3453 CNRS, US7 Inserm, UCBL) facility: CIQLE (a LyMIC member), especially Elisabeth Errazuriz-Cerda for her help in sample preparation and image acquisition for transmission EM. TOSV strain MRS2010-4319501 was obtained from the European Virus Archive Global (EVAg) project that has received funding from the European Union’s Horizon 2020 research and innovation program under grant agreement No 653316. We thank Pierre-Yves Lozach (IVPC UMR754, INRAE) and Philippe Marianneau (ANSES Lyon, France) for kindly providing us with TOSV antibodies and useful discussion. We also thank Karl-Klaus Conzelmann and Richard E. Randall for providing us with cells. Finally, we thank Alessia Armezzani for revising the manuscript, and members of our laboratories for useful suggestions.

## Supporting information captions

**Fig S1.**
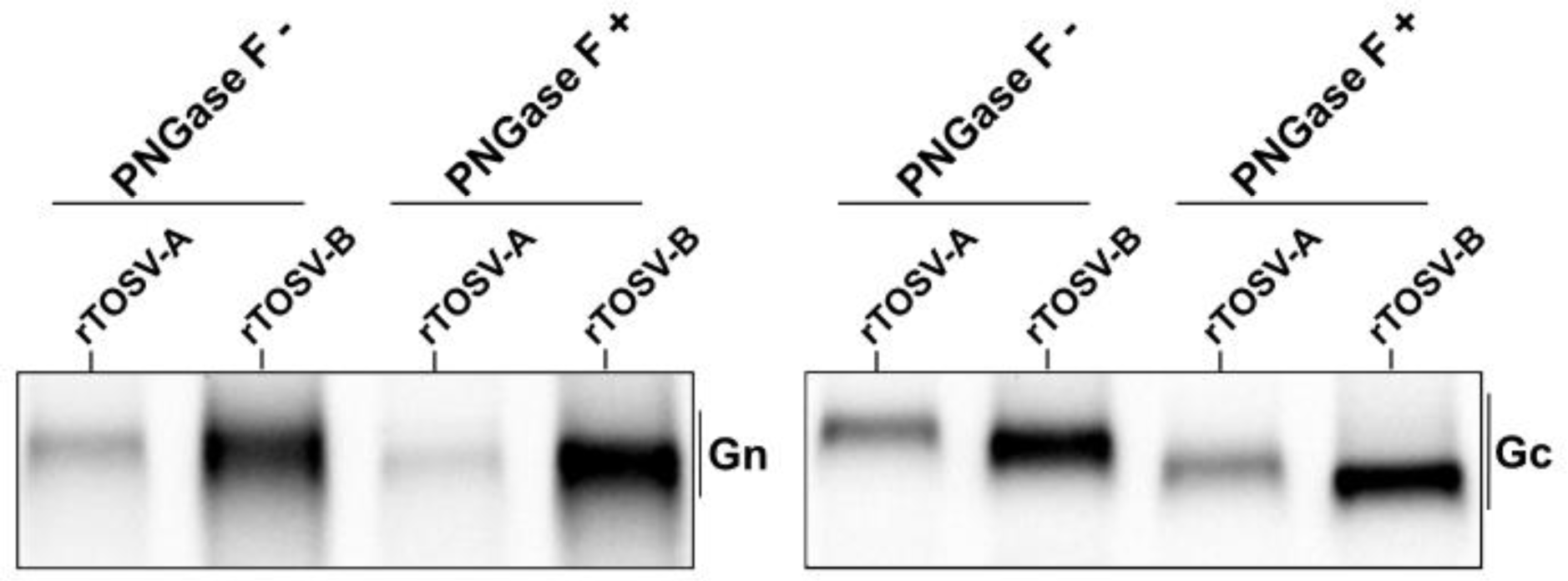
N-glycosylation pattern of rTOSV-A and rTOSV-B Gn and Gc glycoproteins. rTOSV-A and rTOSV-B virions produced from A549 Npro infected cells were purified by ultracentrifugation and treated or not with PNGase-F. The virions were then analysed by western blot with antisera raised against TOSV Gn or Gc proteins as indicated. Note that there is still a difference in the size of rTOSV-A and rTOSV-B Gn glycoproteins even after the PNGase-F treatment.

**Fig S2.**
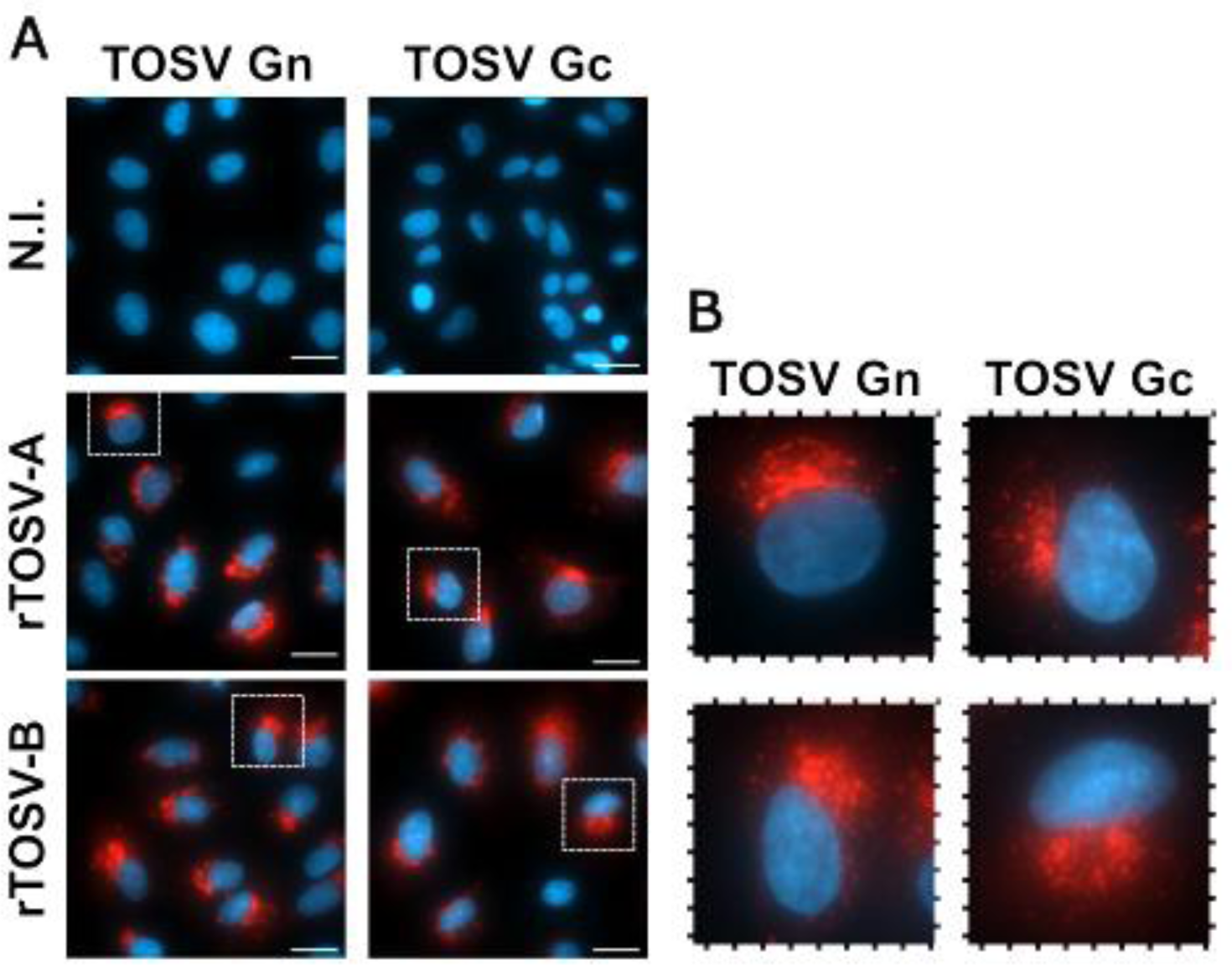
Cellular localization of rTOSV-A and -B Gn and Gc glycoproteins. (A) A549 Npro cells were either non-infected (N.I.) or infected with rTOSV-A or rTOSV-B viruses (MOI=0.3). At 14 hours post-infection, the cells were fixed with PFA and immuno-stained for TOSV glycoproteins with an anti-Gn or anti-Gc antibody followed by a secondary antibody conjugated to Alexa568 (in red). Cell nuclei were labelled with DAPI staining (in blue). Representative pictures of TOSV-infected cells from two independent experiments are shown. The scale bars at the low right corner represent 20 µm. (B) Magnification of areas are indicated with white squares.

## Notes

### Competing Interest Statement

The authors have declared no competing interest.

